# BYSTER: multimodal graph learning resolves bystander versus tumor-specific T cell states in the tumor immune microenvironment

**DOI:** 10.64898/2026.11.24.769066

**Authors:** Haixia Xu, Jianguo Wen, Kexin Huang, Xiaobo Zhou

**Affiliations:** Center for Computational Systems Medicine, McWilliams School of Biomedical Informatics, The University of Texas Health Science Center at Houston, Houston, Texas, 77030, USA; McGovern Medical School, The University of Texas Health Science Center at Houston, Houston, TX 77030, USA; School of Dentistry, The University of Texas Health Science Center at Houston, Houston, TX 77030, USA

**Keywords:** Bystander T cells, Tumor-specific T cells, Tumor microenvironment, T-cell receptor repertoire, Heterogeneous graph neural network, Immunotherapy response

## Abstract

Bystander T cells infiltrate tumors without recognizing tumor antigens and constitute a substantial component of the tumor microenvironment. Whether they are passive residents or active participants in antitumor immunity, and how they influence immunotherapy response, remains poorly understood, largely because transcriptional states and T-cell receptor (TCR) sequences alone do not reliably resolve antigen specificity. Here, we developed BYSTER, a heterogeneous graph neural network that integrates transcriptional representations with paired TCR representations through cross-modal message passing at metacell and metaTCR resolution. Across three datasets with distinct definitions of tumor specificity, BYSTER achieved mean AUROCs of 0.729 to 0.920, outperforming unimodal and conventional multimodal baselines, and ablation analyses confirmed the contribution of cross-modal message passing. Application to lung and pancreatic cancer cohorts revealed distinct biological characteristics of the two predicted populations. Bystander T cells were enriched for naive and memory subsets and MAIT-like cells expressing SLC4A10, and exhibited inflammatory and cellular stress-associated programs, including TNFα signaling via NF-κB and hypoxia. Notably, these transcriptional patterns were consistent across lung and pancreatic cancers, with tumor-specific T cells showing enrichment of cell cycle and proliferation-associated programs in both cancer types. Spatial analyses further revealed distinct inferred communication patterns, with bystander T cells showing associations with inflammatory ligand–receptor interactions involving myeloid populations. Responders to immunotherapy contributed a larger proportion of the predicted bystander compartment, suggesting an association between bystander T-cell abundance and an immune-responsive microenvironment. BYSTER thus links predicted T-cell antigen specificity to transcriptional states, spatial organization and clinical response, highlighting the potential importance of bystander T cells in tumor immunity.

**One-sentence summary:** BYSTER integrates gene expression and TCR data to distinguish predicted bystander and tumor-specific T cells in cancer.

## Introduction

Cancer immunotherapy has transformed the treatment of multiple malignancies by harnessing T-cell mediated antitumor immunity[1]. However, a substantial proportion of patients fail to achieve durable clinical responses[2], , highlighting the need to better understand the cellular mechanisms underlying treatment efficacy. Tumor immune infiltration and the functional states of tumor infiltrating lymphocytes (TILs) are important features of the tumor microenvironment[3, 4]. Nevertheless, T-cell infiltration does not necessarily indicate tumor antigen recognition, as tumors contain substantial populations of bystander T cells that recognize unrelated antigens or lack detectable tumor reactivity. These cells can undergo antigen independent activation in response to inflammatory cytokines and local microenvironmental signals[5, 6].

Accurate identification of bystander and tumor-specific T cells is essential for understanding their distinct contributions to antitumor immunity and immunotherapy response. Previous studies have identified transcriptional markers associated with tumor reactivity, including CD39, PD-1 and CXCL13 [7–9]. However, these markers primarily reflect cellular activation and differentiation states and are not exclusive to tumor-specific T cells. Bystander T cells can also acquire activated, memory or exhausted phenotypes in response to inflammatory cytokines and local microenvironmental signals, resulting in substantial transcriptional overlap between the two populations [10, 11]. T-cell receptor (TCR) sequencing provides complementary information about clonotype identity, clonal expansion and repertoire diversity, but neither TCR sequence nor clonal expansion alone can conclusively determine tumor antigen specificity[12]. Although experimental approaches, including antigen stimulation and functional TCR screening, enable direct assessment of antigen recognition, their scalability remains limited for large clinical cohorts[13]. These challenges highlight the need for computational methods that integrate transcriptional and TCR information to distinguish bystander from tumor-specific T cells and clarify their respective associations with immunotherapy outcomes.

Recent advances in single-cell RNA sequencing (scRNA-seq) and paired TCR sequencing have enabled simultaneous characterization of T-cell transcriptional states and antigen receptor repertoires. Foundation models such as scGPT[14] and SCEPTR[15] provide informative representations of gene expression and TCR sequences, respectively. These developments offer opportunities to integrate complementary molecular information for predicting T-cell antigen specificity. Existing multimodal approaches, however, often rely on independently learned features or their direct concatenation, without explicitly modeling the biological relationships between transcriptional states and TCR identities[16, 17]. T cells sharing the same clonotype can occupy distinct functional states, whereas transcriptionally similar cells may carry different TCRs with distinct antigen recognition properties. Capturing these relationships is particularly important for distinguishing antigen-driven T-cell responses from cytokine-mediated bystander activation. Heterogeneous graph neural networks provide a framework for representing cellular states, clonotype identities and biological associations as interconnected entities, allowing information to propagate across molecular modalities. Such integration may improve the identification of functionally distinct T-cell populations and enable a more comprehensive understanding of the immune features associated with treatment response.

Here, we developed BYSTER, a heterogeneous graph neural network that integrates transcriptional and paired TCR representations to distinguish bystander from tumor-specific T cells. BYSTER combines RNA embeddings with TCR embeddings at metacell and metaTCR resolution, propagating information across transcriptional similarity, clonotype sharing, pathway activity and TCR gene segment relationships through cross-modal message passing. We validated BYSTER across three datasets representing distinct definitions of tumor specificity, including database-inferred antigen matching experimentally determined T-cell specificity, and further applied it to non-small cell lung cancer and pancreatic ductal adenocarcinoma cohorts. By integrating analyses of cellular composition, transcriptional programs, TCR repertoire characteristics, differentiation dynamics, clinical outcomes and spatial interactions, BYSTER provides insights into the distinct roles of bystander and tumor-specific T cells within the tumor microenvironment and their associations with immunotherapy response.

## Materials and Methods

### Paired single RNA-seq and TCR-seq data collection and preprocessing

Paired single-cell RNA-seq and TCR-seq data were collected from five public sources (Supplementary Table 1). Three datasets spanning 23 cancer types formed a pan-cancer corpus for self-supervised pretraining without specificity labels and subsequent benchmarking using database-inferred annotations. Two additional cohorts, comprising non-small-cell lung cancer data from GSE179994[9] and pancreatic ductal adenocarcinoma data from GSE254250[18], provided experimental annotations for cancer-specific fine-tuning and evaluation.

Clonotypes were defined by paired TRA and TRB CDR3 amino acid sequences. Bystander and tumor-specific labels were assigned according to the antigen-specificity or tumor-reactivity evidence associated with each clonotype and propagated to all cells carrying that clonotype within a sample.

For the pan-cancer corpus, Clonotypes were matched against VDJdb[19] and NeoTCR[20] by exact amino acid identity of both CDR3α and CDR3β sequences within the same reference TCR pair. Matched records were curated using their antigen and epitope annotations. Clonotypes matching tumor-antigen-specific or neoantigen-specific entries were labelled tumor-specific, whereas those matching entries with pathogen-derived antigen specificity were labelled bystander.

For the lung cancer cohort, experimental specificity annotations were adopted from Caushi et al.[21]. Clonotypes matching mutation-associated neoantigen-specific TCR sequences identified by the MANAFEST assay were labelled tumor-specific. Clonotypes matching virus-specific sequences identified by ViraFEST were labelled bystander. These annotations provided experimentally established antigen specificities for both classes.

For the PDAC cohort, functional annotations were adopted from Meng et al.[18]. In the source study, TCRα/β pairs from tumor-infiltrating lymphocyte clonotypes prioritized using gene-expression signatures were synthesized and transiently introduced as mRNA into a human T-cell line with low background reactivity. The engineered cells were co-cultured with autologous tumor cells pretreated with IFNγ, and activation was assessed by flow cytometric measurement of TNFα and CD107a. Clonotypes whose TCRs induced detectable activation above mock and non-reactive controls were labelled tumor-specific. Tested clonotypes without detectable reactivity under the same assay conditions were labelled bystander.

### Metacell–metaTCR graph construction

We represented each biological sample as a heterogeneous graph that jointly describes transcriptional states, T-cell receptor (TCR) clonotypes, pathway activity and TCR gene segment usage. Each graph comprised four types of nodes (metacells, metaTCRs, gene sets, and segments) interconnected through 16 types of directed relations. Graph construction used gene expression counts and TCR annotations from cells with paired TCR α and β complementarity-determining region 3 (CDR3) amino acid sequences. Sample identifiers defined graph membership, each sample was assigned to exactly one patient, and cell identifiers linked the expression and TCR layers throughout preprocessing.

### Cell-level transcriptional representations

We computed cell embeddings with a frozen, pretrained scGPT model[14], with gene identifiers obtained from the expression matrix index for both pretraining and fine-tuning. Each cell was represented by a 512-dimensional embedding, which was subsequently normalized to unit Euclidean norm.

### Metacell construction

We constructed metacells independently within each sample using SEACells[22], taking the normalized scGPT embeddings as the kernel-construction representation. For a sample containing Nₛ cells, the requested number of metacells was

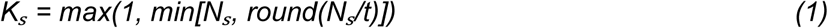

where t denotes the target number of cells per metacell. Samples with Kₛ = 1 were assigned to a single metacell. Random seeds were set to 42 plus the sample index. Hard assignments from the fitted model defined metacell membership, and metacells did not span samples or patients.

### Metacell features

Each metacell node combined an aggregate transcriptional representation with statistics describing its clonotype composition. For metacell m with member cells Cₘ, the normalized scGPT embeddings eᵢ were averaged and re-normalized:

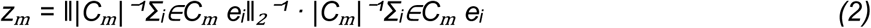

Raw gene expression counts were summed across cells within each metacell to generate pseudobulk expression profiles, which were subsequently used to quantify gene-set activity.

Clonotype composition was summarized from the paired CDR3 sequences of the member cells. Let nₘᶜ denote the number of cells of clonotype c in metacell m, and let pₘᶜ = nₘᶜ/nₘ, where nₘ = Σᶜ nₘᶜ. We quantified clonotype diversity by the Shannon entropy

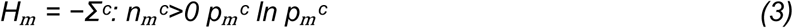

and characterized composition by the clonotype richness Rₘ (the number of distinct clonotypes), the normalized entropy Hₘ/ln Rₘ for Rₘ > 1 (zero otherwise) and the dominant clonotype fraction maxᶜ pₘᶜ. Together with the number of cells carrying paired TCRs, Tₘ, and their fraction among metacell members, fₘ = Tₘ/nₘ, these quantities formed the metacell feature vector

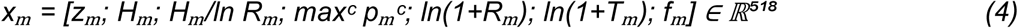

### MetaTCR definition and encoding

MetaTCR nodes represented clonotypes within individual samples, allowing both paired TRA/TRB and single-chain TCR sequences. Cells sharing identical available CDR3 amino acid sequences within the same sample were assigned to the same MetaTCR node, identified by the combination of the sample identifier and available TRA and/or TRB CDR3 sequences. A patient-level clonotype identifier was similarly defined by combining patient identity with the available CDR3 sequences to track clonotypes across samples. Each node stored its constituent cell count and annotations for individual TCR gene segments.

MetaTCR features were generated with a frozen SCEPTR encoder[15]. TRAV, TRAJ, TRBV and TRBJ annotations were standardized with tidytcells[23] to functional human allele-level calls. Nodes with valid annotations for all four fields were encoded with the default SCEPTR model together with their paired CDR3 sequences; nodes with incomplete standardized annotations were encoded with the SCEPTR CDR3-only variant. Unique encoder inputs were embedded into 64-dimensional vectors, which were then assigned to their corresponding nodes.

### Gene-set activity

Each graph contained 50 gene-set nodes corresponding to the MSigDB Hallmark 2020 collection, represented by 50-dimensional one-hot vectors ordered as in the gene-set file. Pathway activity was computed from metacell pseudobulk counts, normalized to 10,000 counts per metacell and transformed with ln(1+x). Gene symbols were matched to Hallmark gene sets case-insensitively, and at least five matched genes were required for each set. For metacell m and gene set h with matched genes Gₕ, the activity score was

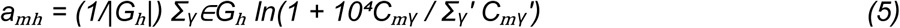

where Cₘᵧ is the pseudobulk count of gene g. Scores were standardized across all metacells of the dataset being processed:

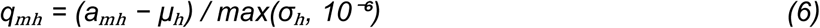

where μₕ and σₕ denote the mean and population standard deviation of the corresponding gene set. Standardization was performed independently for each preprocessing run.

### Segment nodes

TCR gene segment identities were represented by a fixed vocabulary of 171 entries(Supplementary Table 2) shared between pretraining and fine-tuning, covering TRAV, TRAJ, TRBV, TRBD and TRBJ entries, including one unknown category per field. Every graph included the full vocabulary, and each segment node received a one-hot feature vector.

### Graph relations

Edges encoded transcriptional similarity, pathway activity, cell membership, shared TRB sequences and gene segment usage, as follows.

#### Transcriptional similarity

Pairwise cosine similarities were computed between the 512-dimensional aggregate embeddings zₘ. Each metacell was connected by outgoing edges to its k = min(10, Mₛ − 1) most similar other metacells within the sample, where Mₛ is the number of metacells in the sample; neighbor selection therefore defined a directed graph. Similarities on the selected edges were min–max scaled to the interval [0, 1] within the sample,

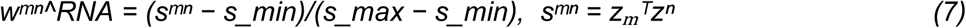

and were set to one when the similarity range was below 10⁻⁸. Samples containing a single metacell had an empty transcriptional-similarity relation.

#### Pathway activity

Every metacell was connected to all 50 gene-set nodes with weight

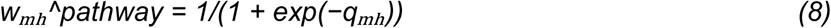

and each gene-set node was connected back to the corresponding metacell with the same weight. The standardized scores qₘₕ, before the sigmoid transformation, were retained as edge attributes and served as regression targets during pretraining. This construction yielded 50Mₛ edges in each direction for a sample containing Mₛ metacells.

#### Metacell–metaTCR membership

An edge connected metacell m to metaTCR c whenever the metacell contained at least one cell assigned to that metaTCR, with weight

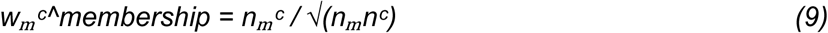

where nᶜ is the total number of cells assigned to metaTCR c within the sample. This normalisation expresses shared membership relative to the sizes of both nodes, retaining the distribution of a receptor identity across transcriptional states and the receptor composition of each state. Each membership edge was accompanied by a reverse edge carrying the same weight under a separate relation type.

#### Shared TRB CDR3

Distinct metaTCR nodes with identical TRB CDR3 amino acid sequences within a sample were connected in both directions, without self-edges. For each shared sequence b, the group abundance was Aᵇ = Σᶜ: CDR3βᶜ=b nᶜ; each edge within the group received the value ln(1 + Aᵇ), which was subsequently min–max scaled to [0, 1] across shared-TRB edges in the sample, with weights set to one when the range was below 10⁻⁸. These edges encode exact β-chain CDR3 sharing without requiring the paired α-chain CDR3 sequences to match; groups containing a single metaTCR node contributed no edges.

#### Gene segment usage

Five relations connected metaTCR nodes to the TRAV, TRAJ, TRBV, TRBD and TRBJ segment nodes, each paired with a reverse relation. Annotations were split on commas, semicolons and vertical bars; allele suffixes were removed, repeated calls deduplicated and ‘/DV’ standardized to ‘DV’. Each resulting gene call generated a unit-weight edge to its vocabulary entry; calls absent from the vocabulary were assigned to the unknown node of the corresponding field, whereas missing annotations yielded no edge.

### Graph schema and supervision labels

The resulting heterogeneous graph comprised 16 directed relation types: one transcriptional similarity relation, two pathway relations, two cell membership relations, one shared TRB relation and ten TCR segment usage relations. Pretraining graphs contained node features, graph connectivity, edge weights and identifier mappings. Fine-tuning graphs additionally included binary labels assigned to MetaTCR nodes. Experimental labels were transferred from constituent cells to their corresponding MetaTCR nodes, requiring consistent labels among cells within each node. All MetaTCR nodes in the fine-tuning graphs were designated as supervised targets.

### Patient level pseudo-bulk differential expression

Raw gene counts were aggregated by patient and predicted label to generate CD8 T-cell pseudobulk profiles. Differential expression analysis was performed using PyDESeq2 with patient identity included as a covariate, retaining patients with at least 10 cells in each predicted group. Low-count genes were excluded. Genes with an adjusted P value < 0.05 and |log₂ fold change| > 1 were considered differentially expressed between tumor-specific and bystander cells.

### Gene set enrichment analysis

Genes were ranked by their Wald statistics and analyzed using preranked GSEA implemented in GSEApy[24] against Hallmark, Gene Ontology Biological Process and KEGG[25] gene sets from MSigDB v2026.1[26]. Gene sets containing 15–500 matched genes were tested using 1,000 permutations, with significance defined as FDR q < 0.05. Positive and negative normalized enrichment scores indicated enrichment in tumor-specific and bystander cells, respectively.

### Comparison of regulon activity

SCENIC regulon AUC scores were averaged by patient and predicted label, retaining patients with at least 10 cells in each group. Regulon activity was compared between tumor-specific and bystander groups using two-sided paired Wilcoxon signed-rank tests with Benjamini–Hochberg correction. Adjusted P values < 0.05 were considered significant. Effect sizes were defined as the median within-patient difference in mean AUC.

### T-cell clonality and association with pathological response

Clonotypes were defined by identical paired TCR α- and β-chain CDR3 amino acid sequences, with expansion defined as at least 3 CD8 T cells per clonotype within a sample. For each patient, cancer type and predicted label, expansion rates were calculated as the proportion of CD8 T cells belonging to expanded clonotypes. Rates were compared between responders and non-responders using two-sided Mann–Whitney U tests in the cohort. Predictive performance was assessed using receiver operating characteristic curves and area under the curve.

### Survival analysis

Patient-level T-cell annotations were integrated with recurrence-free survival (RFS) metadata. For each patient, the proportion of each T-cell subset was calculated separately within the predicted bystander and tumor-specific compartments, using the total number of CD4⁺ and CD8⁺ T cells in the corresponding compartment as the denominator. For each subset, a proportion difference was calculated by subtracting its proportion in the tumor-specific compartment from its proportion in the bystander compartment. Patients were stratified into high- and low-groups using the cohort median. The same denominator definition was used in the combined NSCLC analysis and in separate analyses of LUAD and LUSC. RFS was estimated using the Kaplan–Meier method and compared between groups using the log-rank test.

### Velocity analysis

RNA velocity was estimated from spliced and unspliced counts in CD8⁺ T cells using the dynamical model in scVelo (v0.3.3). After filtering and normalization, neighborhood moments were calculated using 20 principal components and 30 nearest neighbors. The top 200 genes ranked by the product of spliced and unspliced moment variances were selected for dynamical fitting, yielding 172 velocity genes. Bystander and tumor-specific T cells were visualized separately using a shared UMAP embedding and velocity model. To assess differences in cell-state transition preferences, transition weights were restricted to cells within the same sample, with self-transitions removed and rows renormalized.

## Results

### Selected transcriptional and clonal features show limited discrimination of clinical response in the lung cancer cohort

The relationship between T-cell transcriptional states, TCR clonotypes and clinical response was examined using paired single-cell RNA and TCR sequencing data from the GSE179994 lung cancer dataset. Bystander annotations were retained from the original dataset. Cell states, clinical response, bystander labels and the five most abundant clonotypes were visualized on a shared UMAP embedding (Figure 1A). The exhausted T-cell (Tex) lineage was defined as Tex cells and non-Tex cells sharing their TCR clonotypes.

**Figure 1:**
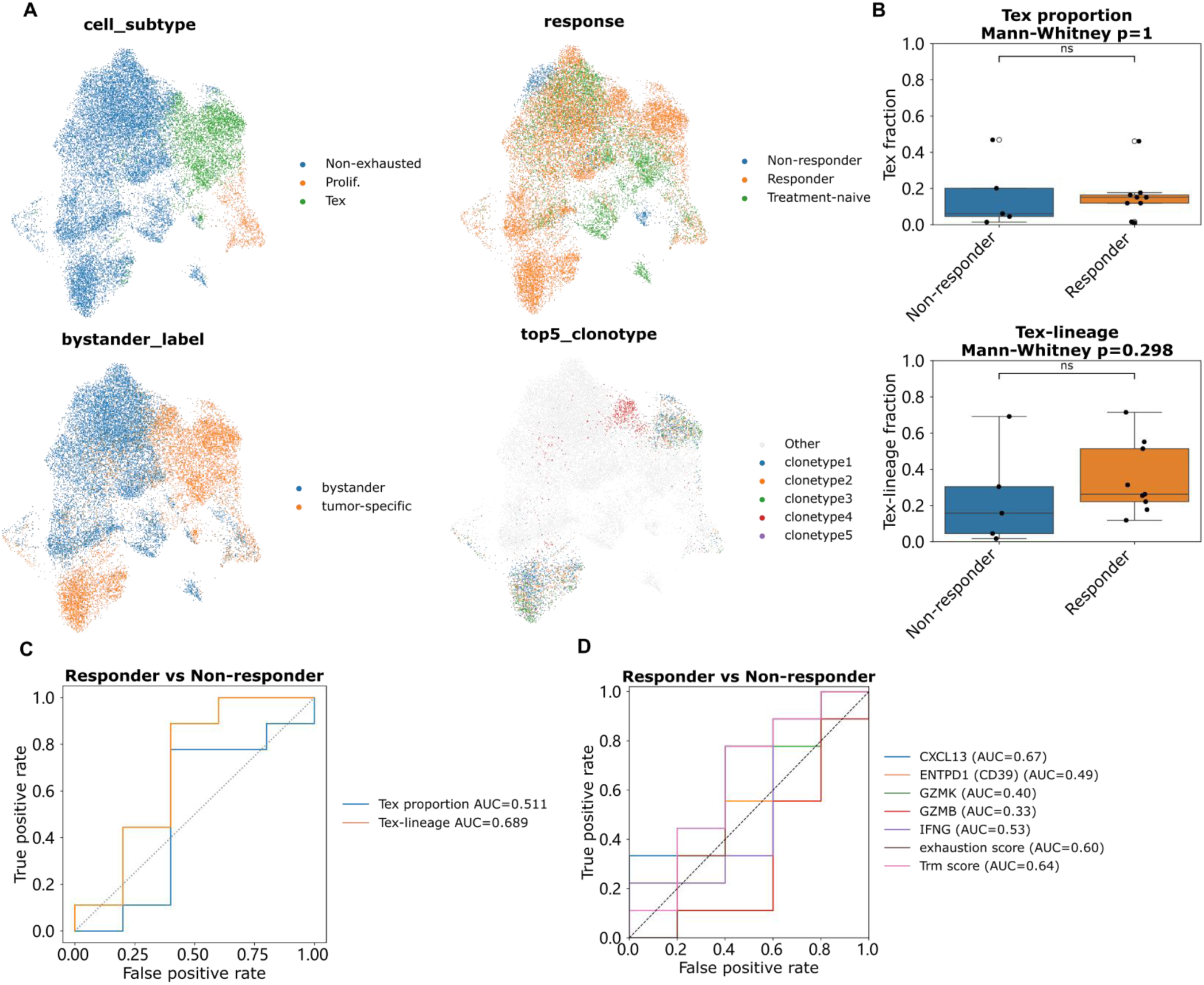
Association of transcriptional features and Tex-lineage abundance with sample-level clinical response in the GSE179994 lung cancer cohort. (A) UMAP embeddings colored by cell subtype, response status, bystander annotation and the five most abundant TCR clonotypes. (B) Comparison of Tex and Tex-lineage fractions between responder and non-responder samples. The Tex lineage comprises Tex cells and non-Tex cells sharing a TCR clonotype with Tex cells. Differences were assessed using the Mann–Whitney U test. ns, not significant. (C) ROC curves assessing sample-level discrimination of responders from non-responders using Tex and Tex-lineage fractions. (D) Sample-level ROC curves for selected literature-derived markers of putative tumor specificity or related T-cell states: CXCL13, ENTPD1 (CD39), GZMK, GZMB and IFNG, together with exhaustion and tissue-resident memory T-cell (Trm) scores. AUROCs are indicated in the legend.

**Supplementary Figure 1:**
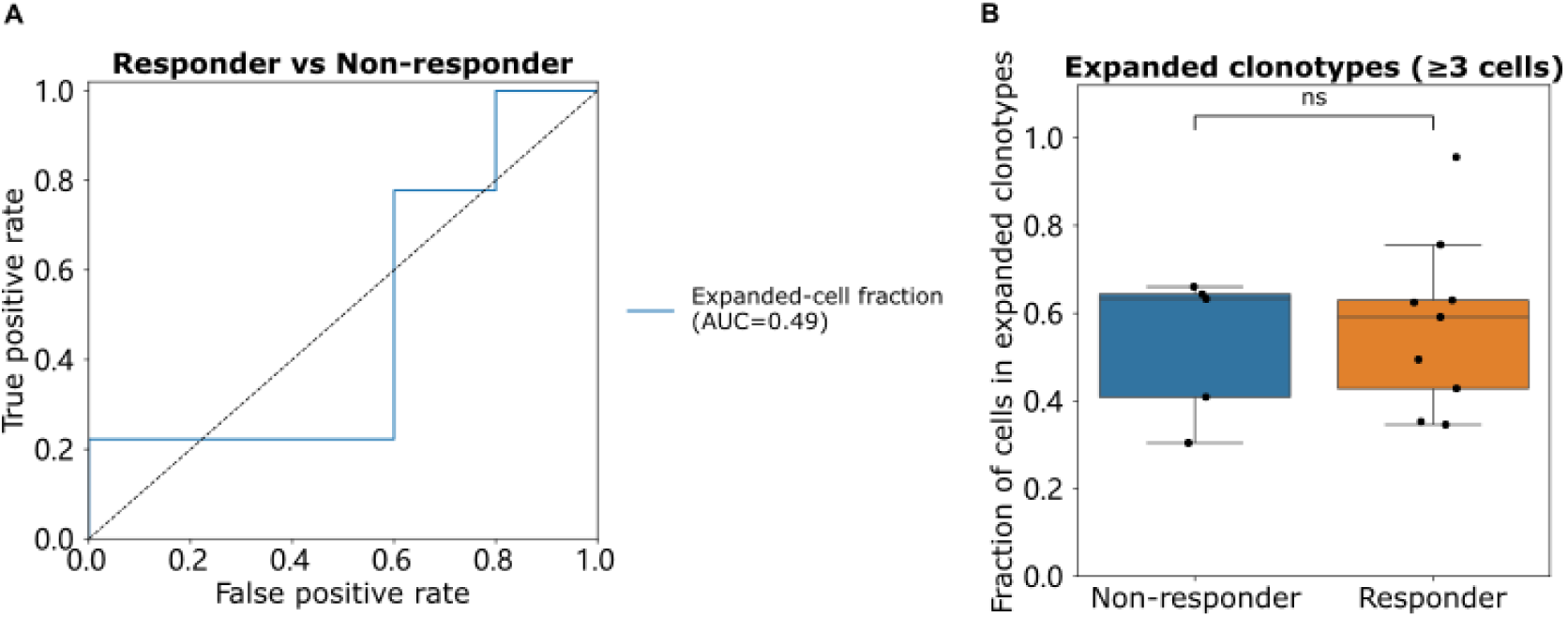
Association of TCR clonal expansion with sample-level clinical response. (A) ROC curve assessing discrimination of responder from non-responder samples using the expanded-cell fraction. The diagonal dashed line indicates chance-level discrimination. (B) Comparison of expanded-cell fractions between responder and non-responder samples. Each point represents a sample. ns, not significant.

Neither Tex nor Tex-lineage fractions differed significantly between responder and non-responder samples (Figure 1B). Tex abundance showed near-chance discrimination of response status (AUROC = 0.511), whereas Tex-lineage abundance provided modest discrimination (Figure 1C). Selected transcriptional markers and signature scores exhibited varying abilities to discriminate clinical response, with CXCL13 showing modest discriminatory performance (Figure 1D).

TCR clonal expansion alone was then evaluated for its association with clinical response. Clonotypes detected in at least three cells were classified as expanded. The proportion of cells belonging to expanded clonotypes showed near-chance discrimination between responder and non-responder samples (Supplementary Figure 1A), with no significant difference between groups (Supplementary Figure 1B), suggesting that individual transcriptional features and clonal expansion metrics offer limited discrimination of sample-level clinical response in this cohort.

### BYSTER integrates transcriptional and TCR information for bystander T-cell classification

The limited ability of transcriptional features or TCR clonal expansion alone to distinguish clinical response motivated a multimodal approach integrating both sources of information. BYSTER was therefore developed as a heterogeneous graph neural network to classify bystander and tumor-specific T cells, enabling assessment of their association with clinical phenotypes (Figure 2A). The framework integrates scGPT-derived RNA representations and SCEPTR-derived paired TCR representations organized into MetaCell and MetaTCR groups. The graph captures transcriptional similarity, shared TCRβ sequences and cell-membership relationships, alongside Hallmark pathway and TCR segment information. Message passing propagates information within and between the RNA and TCR layers. For classification, each updated MetaTCR representation is combined with the context of its connected MetaCells and passed to a prediction head.

**Figure 2:**
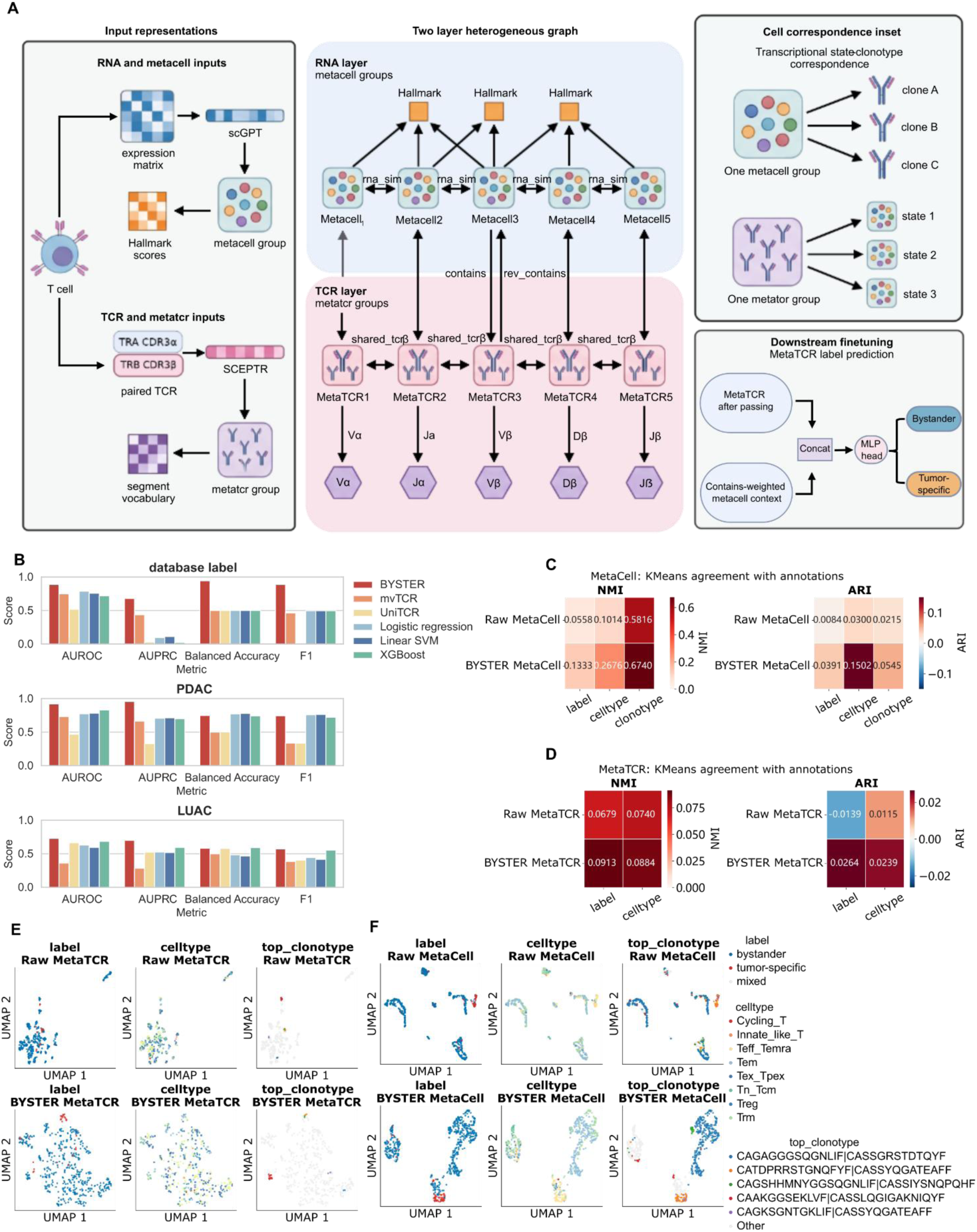
BYSTER framework, fine-tuning performance and representation analysis. (A) Overview of BYSTER. RNA and paired TCR representations generated by scGPT and SCEPTR are grouped into MetaCells and MetaTCRs, respectively. A heterogeneous graph integrates transcriptional similarity, shared TCRβ sequences, cell membership, Hallmark pathways and TCR segment information through message passing. Updated MetaTCR representations are combined with connected MetaCell context for bystander versus tumor-specific T cell classification. The inset illustrates transcriptional state and clonotype correspondence. (B) Fine-tuning performance of BYSTER, mvTCR, UniTCR, logistic regression, SVM and XGBoost on three labelled datasets: the database-label dataset and the experimentally labelled pancreatic ductal adenocarcinoma (PDAC) and lung cancer (LUAC) datasets. Performance is evaluated using area under the receiver operating characteristic curve (AUROC), area under the precision–recall curve (AUPRC), balanced accuracy and F1 score. (C) Agreement between K-means clusters of MetaCell representations and labels, cell-type annotations and clonotype identities in the database-label dataset before (Raw MetaCell) and after (BYSTER MetaCell) message passing. Agreement is quantified using normalized mutual information (NMI) and the adjusted Rand index (ARI). (D) Corresponding analysis of MetaTCR representations before (Raw MetaTCR) and after (BYSTER MetaTCR) message passing, evaluated against labels and cell-type annotations. (E, F) UMAP visualizations of MetaTCR (E) and MetaCell (F) representations before (Raw) and after (BYSTER) message passing. In both panels, top and bottom rows show representations before and after message passing, respectively, with (E) additionally colored by label, cell type and selected clonotypes.

**Supplementary Figure 2:**
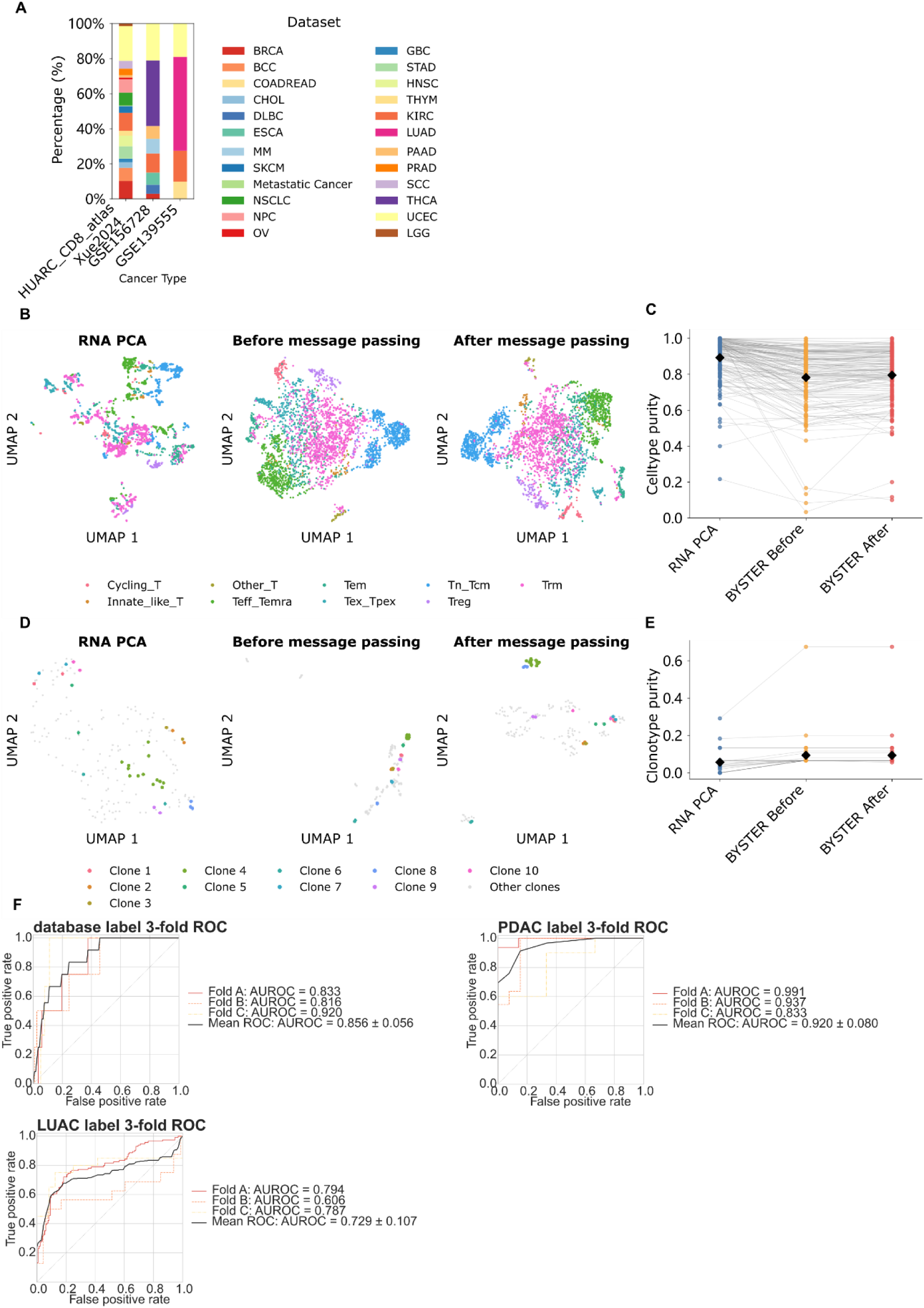
Pretraining data composition, three-fold evaluation and neighborhood structure of BYSTER representations. (A) Cancer-type composition of the three datasets used for pretraining. Stacked bar plots show the relative contribution of each cancer type within each dataset. (B) UMAP visualizations comparing RNA principal component analysis (PCA) representations with BYSTER representations before and after message passing, colored by cell-type annotation. (C) Cell-type neighborhood purity across the three representations in (B), calculated using the 100 nearest neighborhoods. (D) Corresponding UMAP visualizations colored by top ten clonotypes, with remaining clonotypes shown in grey. (E) Clonotype neighborhood purity across the three representations in (D), calculated using the 15 nearest neighborhoods. In (C) and (E), grey lines connect corresponding observations across representations. (F) Receiver operating characteristic (ROC) curves from three-fold evaluation of BYSTER fine-tuning on the database-label, pancreatic ductal adenocarcinoma (PDAC) and lung cancer (LUAC) datasets. Fold-specific AUROCs were 0.833, 0.816 and 0.920 for the database-label dataset; 0.991, 0.937 and 0.833 for PDAC; and 0.794, 0.606 and 0.787 for LUAC. The corresponding mean ± standard deviation AUROCs were 0.856 ± 0.056, 0.920 ± 0.080 and 0.729 ± 0.107, respectively. Black curves indicate mean ROC curves, and diagonal lines indicate chance-level discrimination.

BYSTER was pretrained on three datasets spanning multiple cancer types (Supplementary Figure 2A). Antigen-specificity annotations were assigned to cells in the pretraining corpus by matching TCR CDR3 sequences against VDJdb and NeoTCR. Subsequent fine-tuning on the database-annotated dataset achieved a mean AUROC of 0.856 ± 0.056 across three folds (mean ± standard deviation; Supplementary Figure 2F). Corresponding three-fold evaluations on the experimentally labelled PDAC and LUAC datasets yielded mean AUROCs of 0.920 ± 0.080 and 0.729 ± 0.107, respectively (Supplementary Figure 2F).

To characterize the pretrained representations, we compared RNA PCA with BYSTER embeddings before and after message passing. UMAP visualizations showed changes in the organization of cell types and highlighted clonotypes across these representations (Supplementary Figure 2B, D). Neighborhood analyses quantified these changes using cell-type purity at 100 nearest neighborhood and clonotype purity at 15 nearest neighborhood (Supplementary Figure 2C,E). BYSTER representations showed higher clonotype purity than RNA PCA and increased after message passing, indicating that the pretrained representations captured additional clonotype structure. We next compared BYSTER with mvTCR, UniTCR, logistic regression, linear support vector machine (SVM)[27] and XGBoost[28] after fine-tuning on the database-label dataset and the experimentally labelled PDAC and LUAC datasets (Figure 2B). BYSTER achieved the highest AUROC on the database-label and LUAC datasets and the highest AUPRC on the PDAC dataset.

The effects of message passing on the learned representations were further evaluated in the database-annotated dataset. Message passing consistently improved the agreement of MetaCell K-means clusters with cell-type annotations, clonotype identities and antigen-specificity labels (Figure 2C), withthe largest absolute gain observed for cell-type annotations. MetaTCR representations showed improvements for antigen-specificity and cell-type agreement (Figure 2D). UMAP visualizations illustrated the corresponding changes in both representation spaces (Figure 2E,F).

Together, these results suggest that BYSTER’s advantage arises from its heterogeneous graph architecture rather than feature richness alone. By enabling information exchange between transcriptional states and TCR sequences during representation learning, BYSTER achieved strong classification performance across three datasets with distinct label sources, outperforming both unimodal and conventional multimodal baselines. Message-passing ablation analyses further demonstrated the contribution of cross-modal information integration, showing that message passing improved the alignment of learned representations with cell states, clonotype identities and antigen specificity beyond simple feature concatenation.

### BYSTER identifies distinct cellular compositions and transcriptional programs in bystander and tumor-specific T cells in NSCLC

To characterize the cellular composition and transcriptional programs of bystander and tumor-specific T cells, we fine-tuned BYSTER on experimentally labelled lung cancer data and then applied the resulting model to an independent NSCLC cohort[29] comprising LUAD and LUSC samples. The two populations occupied partially overlapping regions of the transcriptional landscape but differed in cellular composition (Figure 3A–C). Bystander T cells were enriched for CD4⁺ naive and memory subsets, whereas tumor-specific T cells contained higher proportions of CD8⁺ effector memory (Tem), tissue-resident memory (Trm) and exhausted (Tex) cells. Within the CD8⁺ compartment, bystander cells contained higher proportions of memory T (Tm) and mucosal-associated invariant T cells, whereas tumor-specific cells were dominated by Tem, Trm and Tex subsets (Figure 3D,E). These compositional differences were consistent across LUAD and LUSC, suggesting a shared pattern across the two histological subtypes (Supplementary Figure 3B-E).

**Figure 3.**
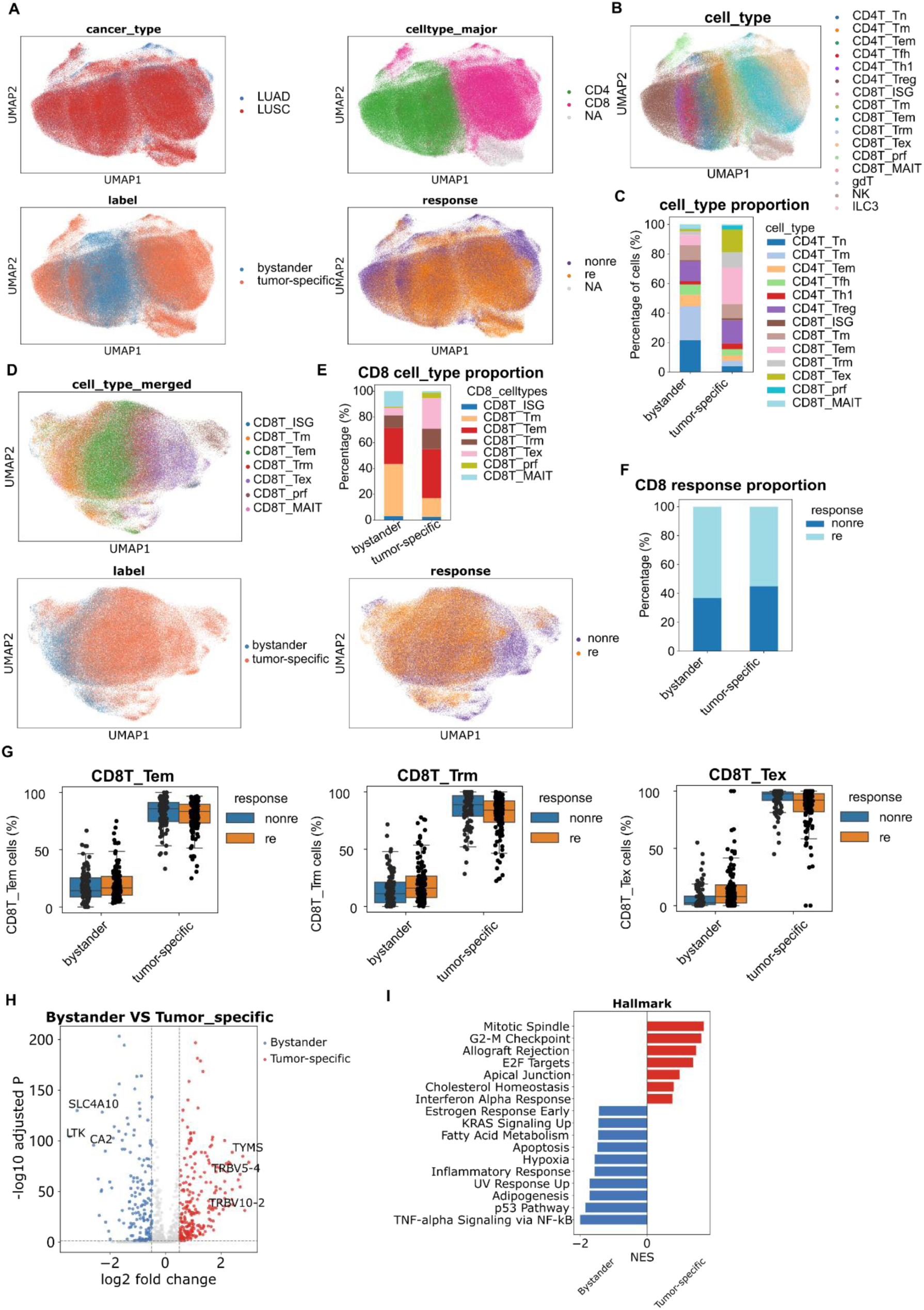
Distinct transcriptional programs of bystander and tumor-specific T cells in NSCLC. (A) UMAP of all T cells from LUAD and LUSC tumors, colored by cancer type, major cell type, BYSTER model-predicted label (bystander vs. tumor-specific), and clinical response. (B) UMAP as in (A), colored by minor T cell subset. (C) Relative proportions of minor T cell subsets within the bystander and tumor-specific compartments. (D) UMAP of CD8⁺ T cells, colored by subset, BYSTER-predicted label, and clinical response. (E) Relative proportions of CD8⁺ T cell subsets within the bystander and tumor-specific compartments. (F) Proportions of CD8⁺ T cells originating from responders and non-responders within each predicted compartment (bystander or tumor-specific). (G) Proportions of predicted bystander and tumor-specific cells within the CD8⁺ effector memory (Tem), tissue-resident memory (Trm), and exhausted (Tex) subsets, compared between responders and non-responders. (H) Volcano plot of differentially expressed genes between bystander and tumor-specific CD8⁺ T cells. (I) Hallmark gene set enrichment analysis comparing predicted bystander and tumor-specific CD8⁺ T cells. Normalized enrichment scores (NES) indicate enrichment in bystander cells (blue, negative NES) or tumor-specific cells (red, positive NES).

**Supplementary Figure 3.**
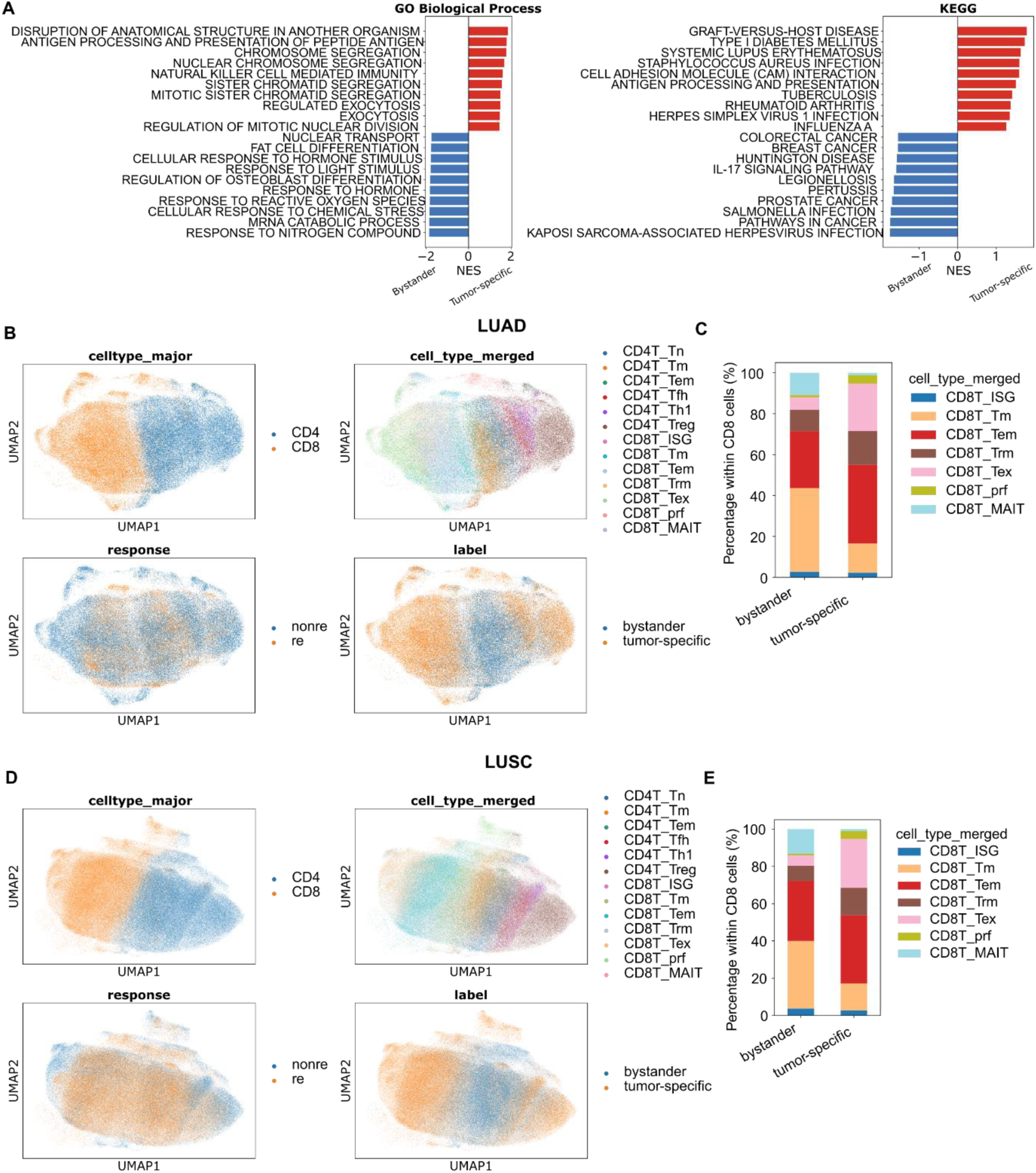
Functional enrichment and cellular composition of bystander and tumor-specific T cells in NSCLC. (A) Gene set enrichment analysis comparing predicted bystander and tumor-specific CD8⁺ T cells using Gene Ontology Biological Process (GO BP) and Kyoto Encyclopedia of Genes and Genomes (KEGG) pathways. Negative normalized enrichment scores (NES, blue) indicate enrichment in bystander cells, and positive NES (red) indicate enrichment in tumor-specific cells. (B) UMAP embeddings of T cells from LUAD samples, colored by major cell type, minor T-cell subset, clinical response, and BYSTER-predicted label. (C) Relative proportions of CD8⁺ T-cell subsets within the predicted bystander and tumor-specific compartments in LUAD. (D) UMAP embeddings of T cells from LUSC samples, colored by major cell type, minor T-cell subset, clinical response, and BYSTER-predicted label. (E) Relative proportions of CD8⁺ T-cell subsets within the predicted bystander and tumor-specific compartments in LUSC.

The distribution of bystander and tumor-specific T cells was then assessed across clinical response groups. Cells from responders accounted for a larger proportion of the bystander compartment than of the tumor-specific compartment (Figure 3F). Although tumor-specific predictions predominated within Tem, Trm and Tex subsets in both response groups, responders generally showed higher median bystander fractions and correspondingly lower tumor-specific fractions within these subsets (Figure 3G). These patterns suggest an association between predicted T-cell identity and clinical response, both across the overall CD8⁺ population and within defined transcriptional states.

Differential expression analysis revealed distinct molecular features of the two CD8⁺ compartments (Figure 3H). Bystander cells showed higher expression of SLC4A10, LTK and CA2, whereas tumor-specific cells showed higher expression of TYMS and TCR variable-region genes, including TRBV5-4 and TRBV10-2. These expression differences accompanied broader divergence in biological programs. Hallmark gene set enrichment analysis revealed enrichment of TNFα signaling via NF-κB, inflammatory response, hypoxia and p53 pathway in bystander cells, whereas tumor-specific cells showed enrichment of G2M checkpoint, E2F targets, mitotic spindle pathways (Figure 3I). Consistently, GO Biological Process analysis identified enrichment of chromosome segregation and antigen processing and presentation in tumor-specific cells, whereas bystander cells showed enrichment of stress- and stimulus-response processes and mRNA catabolism (Supplementary Figure 3A). KEGG analysis further highlighted immune-related pathways in tumor-specific cells, including antigen processing and presentation, while bystander cells showed enrichment of IL-17 signaling and cellular senescence pathways.

These findings highlight distinct cellular compositions and transcriptional programs between the two predicted T-cell populations in NSCLC. Bystander T cells combined a memory-skewed composition with inflammatory and environmental stress-response programs and were over-represented in clinical responders, whereas tumor-specific cells were dominated by proliferating CD8⁺ effector states. The inflammatory, non-proliferative character of the bystander compartment is consistent with cytokine-mediated bystander activation rather than antigen-driven clonal expansion, identifying it as a distinct, response-associated component of the tumor immune microenvironment.

### Bystander and tumor-specific T cells display distinct TCR repertoire features in NSCLC

Given their distinct transcriptional profiles, predicted bystander and tumor-specific T cells were further compared in terms of regulon activity and TCR repertoire features within the NSCLC cohort. SCENIC analysis showed higher activity of JUN family and NFKB1 regulons in bystander cells, consistent with their enrichment of inflammatory and stress response programs. Tumor-specific cells showed higher FOXM1 regulon activity, consistent with their enrichment of cell cycle and mitotic programs (Figure 4A). These populations also differed in TCR segment usage (Figure 4B). Among the displayed TRBV segments, TRBV20-1 was more prominent in bystander cells than in tumor-specific cells, whereas tumor-specific cells showed a more even distribution across their leading TRBV segments. Differences extended to the α-chain repertoire. TRAV13-1 and TRAJ33 were the most abundant displayed segments in bystander cells, whereas TRAV29/DV5 and TRAJ49 predominated in tumor-specific cells. TRBJ usage also differed, with TRBJ2-7 predominating in bystander cells and TRBJ2-1 in tumor-specific cells.

**Figure 4.**
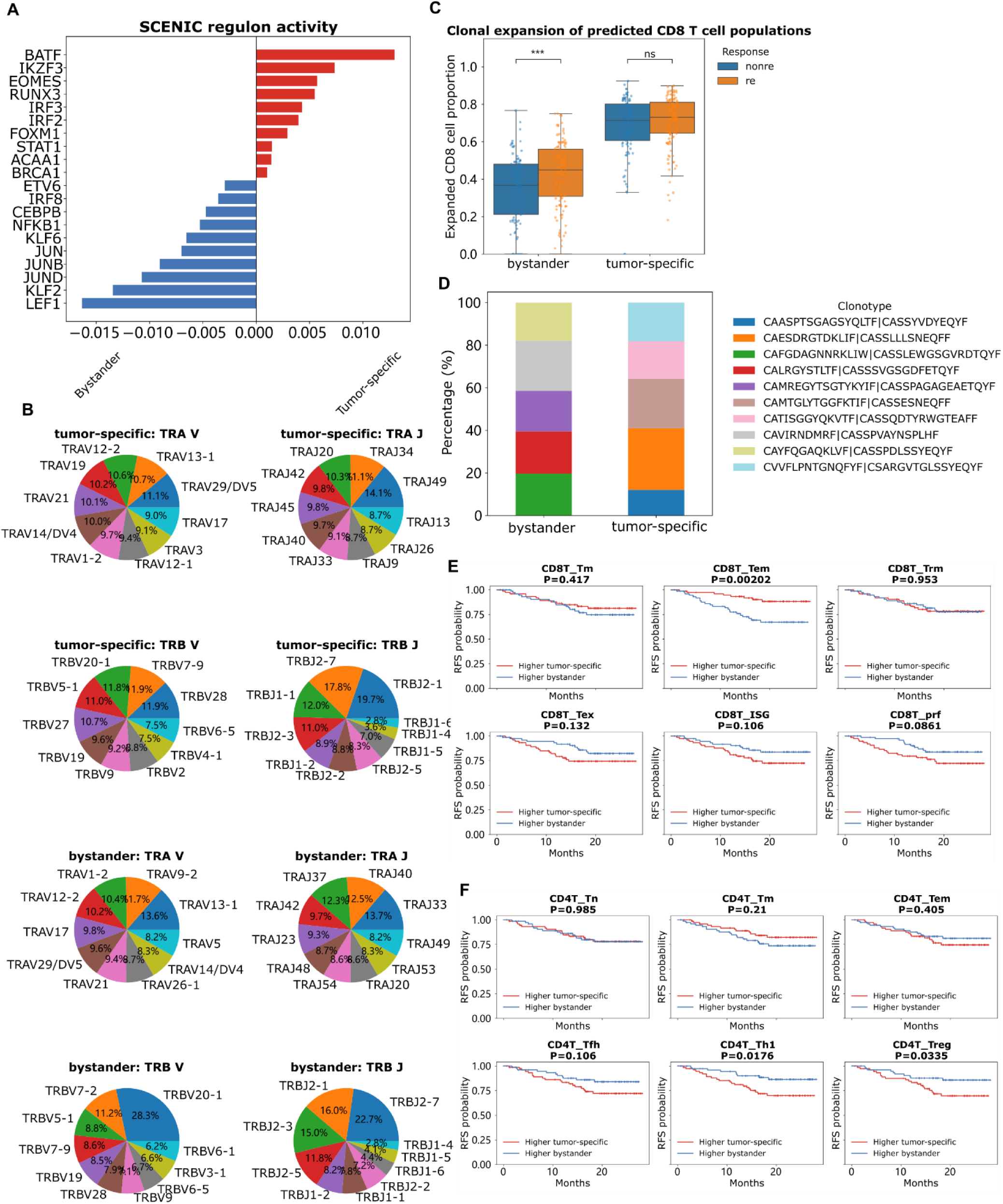
Distinct TCR repertoire features and clinical associations of bystander and tumor-specific T cells in NSCLC. (A) Differential SCENIC regulon activity between BYSTER-bystander and tumor-specific T cells. Blue bars indicate regulons preferentially active in bystander cells, and red bars indicate regulons preferentially active in tumor-specific cells. (B) Relative usage of the ten most abundant V and J segments of the TCR α and β chains in each predicted compartment. Percentages indicate the relative contributions of the displayed segments within each pie chart. (C) Fractions of CD8⁺ T cells belonging to expanded clonotypes in the bystander and tumor-specific compartments, compared between responders (re) and non-responders (nonre). Clonotypes detected in at least three cells were classified as expanded. Statistical significance annotations are shown above the comparisons; ns, not significant. (D) Relative abundance of the five most abundant clonotypes in each predicted compartment. Clonotypes are represented by paired TCR α and β CDR3 sequences, separated by a vertical bar. The two sets of top five clonotypes do not overlap. Each bar is normalized to the total abundance of its five displayed clonotypes. (E) Kaplan–Meier curves of recurrence-free survival (RFS) for patients stratified by compartment-specific proportion differences for the indicated CD8⁺ T-cell subsets: Tm, Tem, Trm, Tex, interferon-stimulated gene-expressing (ISG) and proliferating (prf) cells. (F) Corresponding analyses for CD4⁺ T-cell subsets: Tn, Tm, Tem, Tfh, Th1 and Treg. In (E) and (F), each subset proportion was calculated using all CD4⁺ and CD8⁺ T cells within the corresponding predicted compartment as the denominator.

**Supplementary Figure 4.**
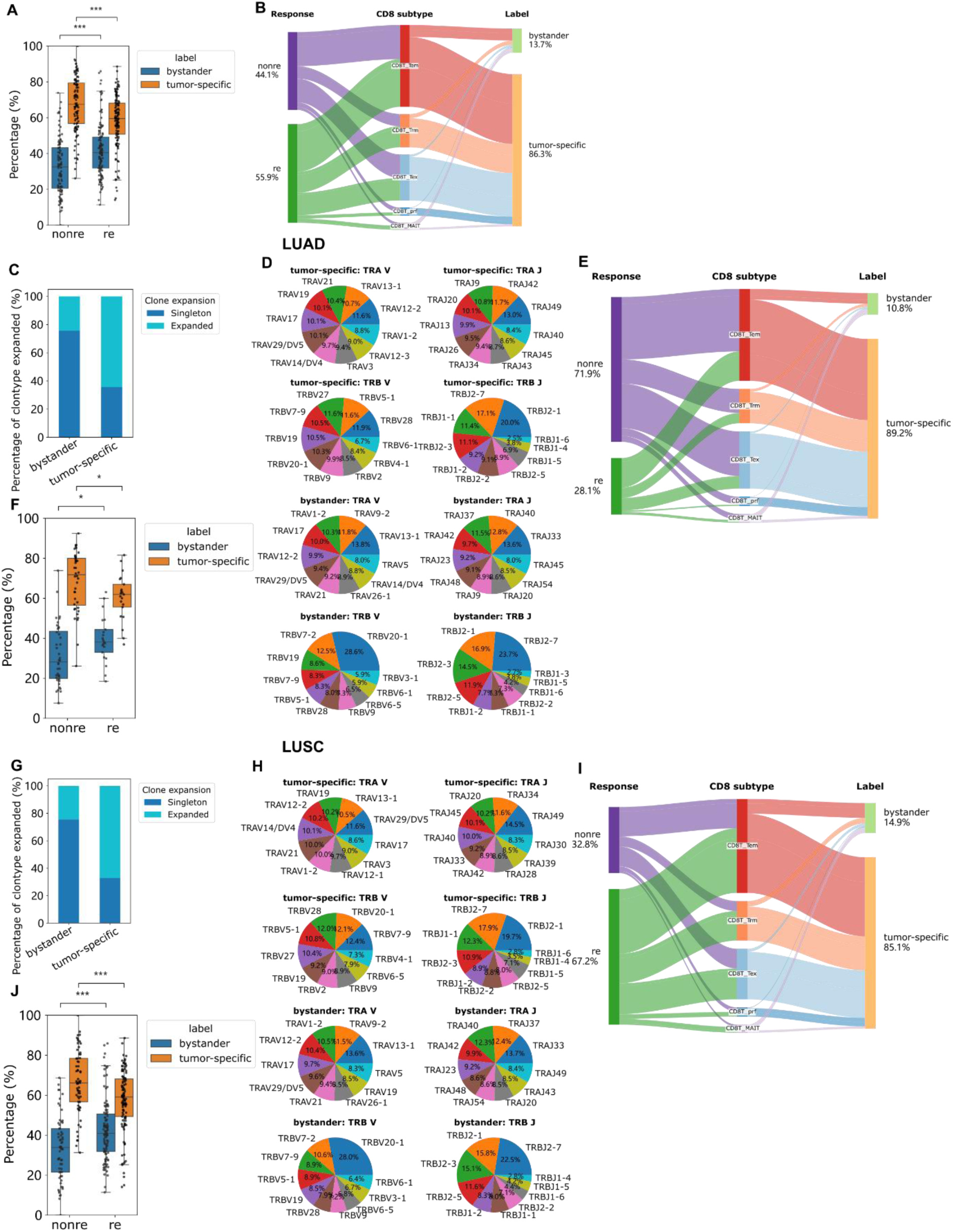
Patient-level proportions by clinical response, CD8⁺ T cell subset distributions, and TCR repertoire features of predicted bystander and tumor-specific T cells in NSCLC. (A) Box plots of patient-level proportions of predicted bystander and tumor-specific T cells by clinical response group in the combined NSCLC cohort. Each dot represents one patient. (B) Sankey diagram linking clinical response groups, CD8⁺ T cell subsets, and BYSTER-predicted labels in the combined NSCLC cohort. (C) Proportions of singleton and expanded clonotypes within the predicted bystander and tumor-specific compartments in LUAD. (D) Usage of the ten most abundant V and J segments for each TCR α and β chain within each predicted compartment in LUAD. Percentages are normalized to the segments displayed in each pie chart. (E) Sankey diagram linking clinical response groups, CD8⁺ T cell subsets, and BYSTER-predicted labels in LUAD. (F) Box plots of patient-level proportions of predicted bystander and tumor-specific T cells by clinical response group in LUAD, displayed as in (A). (G) Proportions of singleton and expanded clonotypes within the predicted bystander and tumor-specific compartments in LUSC. (H) TCR V and J segment usage in LUSC, displayed as in (D). (I) Sankey diagram linking clinical response groups, CD8⁺ T cell subsets, and BYSTER-predicted labels in LUSC. (J) Box plots of patient-level proportions of predicted bystander and tumor-specific T cells by clinical response group in LUSC, displayed as in (A).

Clonal expansion was assessed by defining expanded clonotypes as those detected in at least 3 cells. Tumor-specific CD8⁺ T cells had higher expanded-cell fractions than bystander CD8⁺ T cells in both responders and non-responders (Figure 4C). Within the bystander compartment, this fraction was higher in responders, while the difference between response groups in the tumor-specific compartment was not significant. The tumor-specific compartment also showed a larger expanded fraction in the separate LUAD and LUSC analyses (Supplementary Figure 4C, G). The five most abundant clonotypes in each predicted compartment formed distinct sets of paired TCR α and β CDR3 sequences (Figure 4D). To examine these patterns at the patient and subset levels, we compared the proportions of predicted bystander and tumor-specific T cells across clinical response groups in the combined NSCLC cohort and separately in LUAD and LUSC (Supplementary Figure 4A, F, J). Sankey diagrams showed how clinical response groups and predicted identities were distributed across CD8⁺ T cell subsets in each analysis (Supplementary Figure 4B, E, I)

The association between T-cell subset composition and recurrence-free survival (RFS) was then evaluated. A lower bystander-minus-tumor-specific difference in CD8⁺ Tem proportions was associated with more favorable RFS (Figure 4E). In contrast, higher bystander proportions among CD4⁺ Th1 and Treg cells were associated with more favorable RFS (Figure 4F). These findings suggest that the clinical relevance of predicted T-cell identity varies across transcriptionally defined subsets. These analyses reveal distinct TCR segment usage, dominant clonotypes, and clonal expansion patterns in bystander and tumor-specific T cells.

### BYSTER reveals distinct cellular compositions, clonal architectures and transcriptional programs of bystander and tumor-specific T cells in PDAC

To assess its applicability beyond NSCLC, BYSTER was fine-tuned on experimentally labelled PDAC data, achieving a mean AUROC of 0.920 ± 0.080 across three folds (mean ± standard deviation; Supplementary Figure 2F). BYSTER predictions were based on experimentally defined tumor reactivity, distinguishing bystander from tumor-specific T cells. Predicted identities were mapped onto the PDAC transcriptional landscape alongside experimental condition, major cell type and fine-grained T-cell subset annotations, revealing that both populations spanned multiple transcriptional regions (Figure 5A).

**Figure 5.**
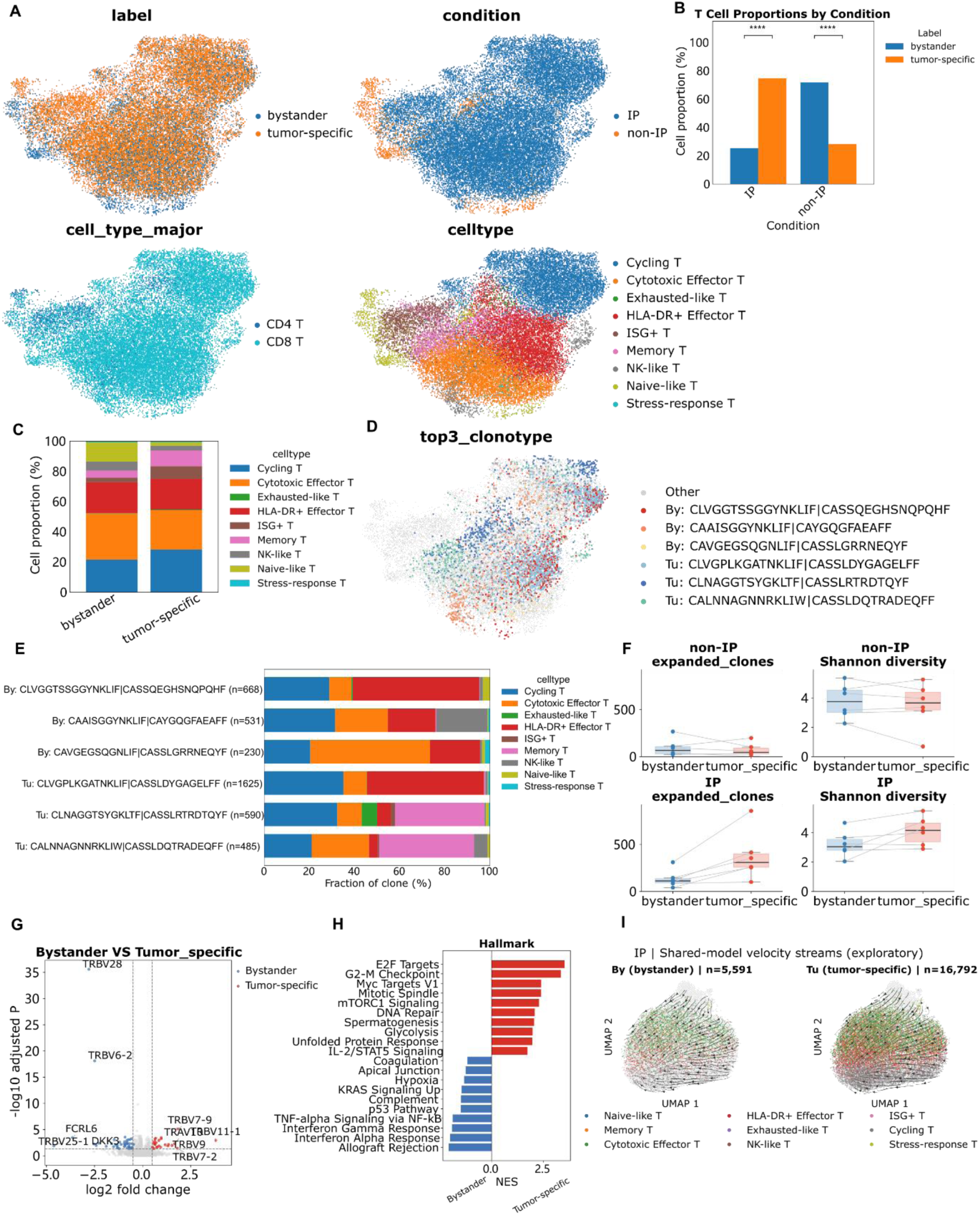
Distinct cellular compositions, clonal architectures and inferred differentiation trajectories of bystander and tumor-specific T cells in PDAC. (A) UMAP visualizations of T cells from pancreatic ductal adenocarcinoma (PDAC) samples, colored by BYSTER-predicted label (bystander versus tumor-specific), experimental condition (IP versus non-IP), major cell type and fine-grained T-cell subset annotation. (B) Proportions of predicted bystander and tumor-specific T cells under IP and non-IP conditions. Brackets indicate comparisons between the two predicted populations within each condition. (C) Relative proportions of annotated T-cell subsets within the predicted bystander and tumor-specific compartments. (D) UMAP visualization highlighting the three most abundant clonotypes within each predicted compartment. Clonotypes are defined by paired TCR α- and β-chain CDR3 amino acid sequences, separated by a vertical bar. Remaining clonotypes are shown in grey. (E) Cell-type composition of the six selected clonotypes shown in (D), comprising the three most abundant clonotypes from each predicted compartment. Stacked bars indicate the relative proportions of annotated T-cell subsets within each clonotype, with the corresponding cell counts indicated. (F) Comparison of expanded clone counts and Shannon diversity between predicted bystander and tumor-specific T-cell compartments, shown separately for non-IP and IP samples. Grey lines connect paired observations. (G) Volcano plot showing differentially expressed genes between predicted bystander and tumor-specific T cells. Positive and negative log2 fold changes indicate higher expression in tumor-specific and bystander T cells, respectively. (H) Hallmark gene set enrichment analysis comparing predicted bystander and tumor-specific T cells. NES indicate enrichment in bystander cells (blue, negative NES) or tumor-specific cells (red, positive NES). (I) RNA velocity streams of CD8⁺ T cells from IP patients, computed with a shared velocity model on all IP cells and visualized separately for bystander (left) and tumor-specific (right) cells. Colored dots, cells of the indicated category; black arrows, smoothed velocity field. n, number of cells of the indicated category.

**Supplementary Figure 5.**
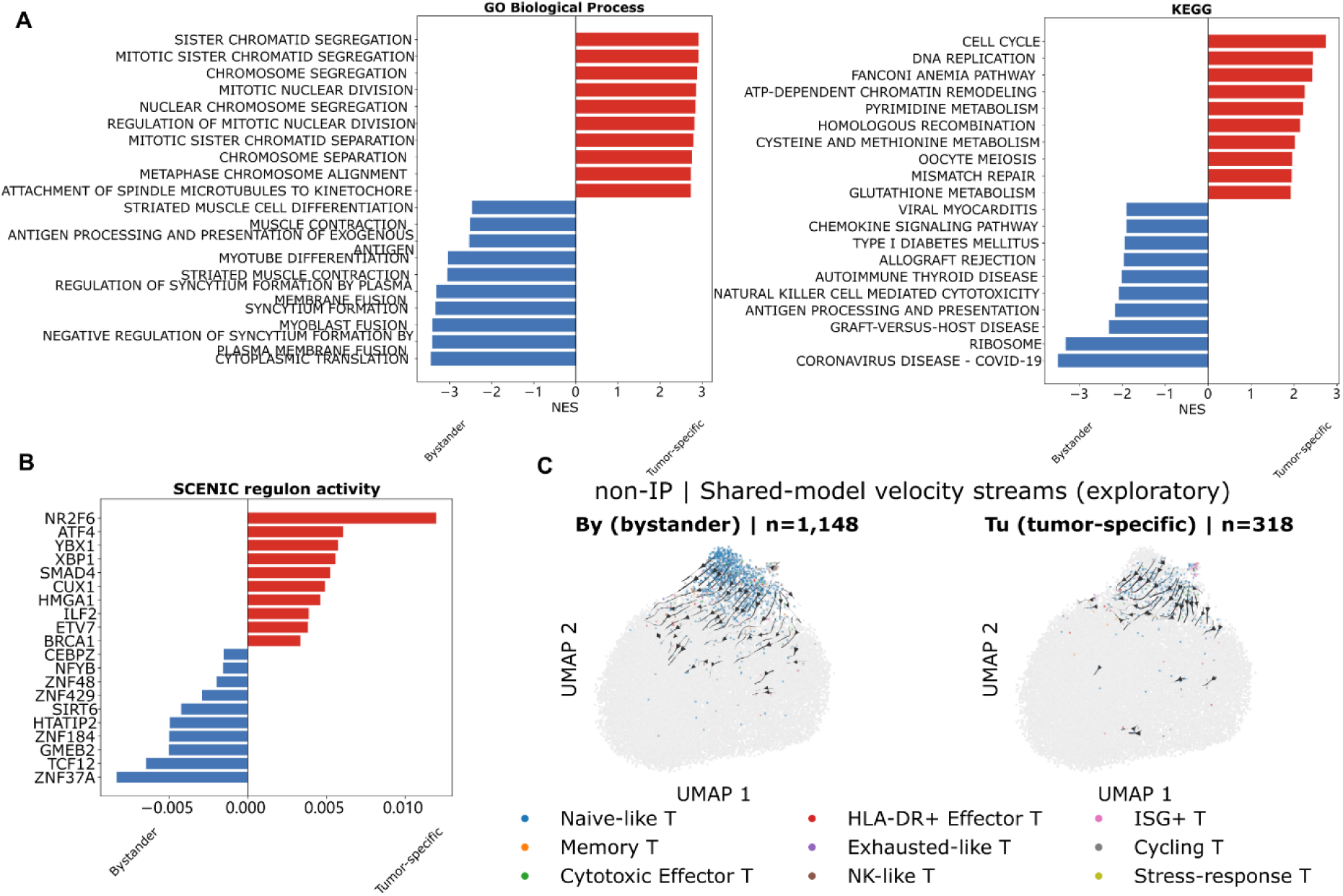
Pathway enrichment and inferred differentiation dynamics of bystander and tumor-specific T cells in PDAC. (A) Gene set enrichment analysis comparing predicted bystander and tumor-specific T cells using GO and KEGG pathway. NES indicate enrichment in bystander cells (blue, negative NES) or tumor-specific cells (red, positive NES). (B) Comparison of SCENIC regulon activities between predicted bystander and tumor-specific T cells. Positive values (red) indicate higher activity in tumor-specific cells, whereas negative values (blue) indicate higher activity in bystander cells. (C) RNA velocity streams of T cells from non-IP samples, computed using a shared velocity model and visualized separately for bystander (left, n = 1,148) and tumor-specific (right, n = 318) cells. Colored dots indicate annotated T-cell subsets, and black arrows represent the inferred velocity field.

The relative abundance of predicted bystander and tumor-specific T cells differed between IP and non-IP conditions (Figure 5B). Predicted tumor-specific cells accounted for approximately 75% of T cells under IP conditions, compared with approximately 28% under non-IP conditions. Fine-grained annotation revealed distinct T-cell subset compositions between the two predicted populations, with differences in the relative abundance of CD4⁺ and CD8⁺ T-cell states (Figure 5A,C). These findings indicate that the two predicted populations shared a broad transcriptional landscape but differed in their relative abundance across experimental conditions and in their T-cell subset compositions. We next investigated whether these compositional differences were accompanied by distinct TCR clonotype distributions and repertoire characteristics. The three most abundant clonotypes from each predicted compartment, defined by paired TCR α-and β-chain CDR3 sequences, showed distinct distributions across the transcriptional landscape (Figure 5D). Examination of these six representative clonotypes further revealed differences in their constituent T-cell subset compositions, indicating heterogeneity in the transcriptional states associated with individual clonotypes (Figure 5E). Repertoire-level analysis further compared expanded clone counts and Shannon diversity between bystander and tumor-specific T cells under IP and non-IP conditions (Figure 5F). Under IP conditions, tumor-specific T cells showed higher expanded clone counts than bystander T cells, suggesting greater clonal expansion in the tumor-specific compartment. In contrast, differences in expanded clone counts were less pronounced under non-IP conditions. Shannon diversity also varied between the two predicted populations and experimental conditions, reflecting differences in TCR repertoire diversity (Figure 5F). T hese findings suggest that differences in T-cell composition under IP conditions are accompanied by distinct patterns of clonal expansion and repertoire diversity between bystander and tumor-specific T cells.

To investigate the molecular differences between the two predicted populations, we performed differential gene expression analysis, which revealed distinct transcriptional profiles in bystander and tumor-specific T cells (Figure 5G). Hallmark gene set enrichment analysis further identified distinct biological programs, with tumor-specific T cells showing enrichment of cell cycle and proliferation-associated pathways, including E2F targets, G2M checkpoint and mitotic spindle, whereas bystander T cells exhibited enrichment of inflammatory and stress-associated pathways, including TNFα signaling via NF-κB, hypoxia and interferon responses (Figure 5H). Notably, the enrichment of cell cycle-related pathways in tumor-specific T cells and inflammatory and stress-associated pathways in bystander T cells was consistent with the patterns observed in NSCLC (Figure 3I), supporting the reproducibility of these distinct transcriptional programs across cancer types. Pathway enrichment and regulon analyses further supported the functional differences between the two populations (Supplementary Figure 5A–B).

We next performed RNA velocity analysis to explore the inferred differentiation dynamics of bystander and tumor-specific T cells. RNA velocity streams of CD8⁺ T cells from IP samples showed differences in the inferred transcriptional dynamics between the two predicted populations (Figure 5I). Additional RNA velocity visualizations further illustrated the distinct patterns of inferred state transitions in bystander and tumor-specific T cells (Supplementary Figure 5C).

Collectively, these analyses reveal distinct cellular compositions, clonal architectures, transcriptional programs and inferred differentiation dynamics between bystander and tumor-specific T cells in PDAC. Bystander T cells showed relatively limited clonal expansion and enrichment of inflammatory and stress-associated pathways, whereas tumor-specific T cells exhibited greater clonal expansion under IP conditions and enrichment of cell cycle and proliferation-associated programs. The concordant transcriptional patterns observed in NSCLC provide further cross-cancer support for these distinct biological characteristics of bystander and tumor-specific T cells within the tumor microenvironment.

### Spatial profiling reveals distinct inferred interaction patterns of bystander and tumor-specific T cells in lung cancer

To place the transcriptional and clonal differences between bystander and tumor-specific T cells in a spatial context, we applied the lung cancer-fine-tuned BYSTER model to a lung cancer Slide-seq dataset [10]. Spatial annotation resolved immune and tumor cell populations across fibrostromal regions, tumor regions and tertiary lymphoid structures (TLS), enabling predicted identities to be mapped alongside T-cell subtypes (Figure 6A). Both predicted populations were broadly distributed across the tissue. Clonotype-resolved mapping showed that most clonotypes had the majority of their cells in tumor regions, followed by fibrostromal regions, with fewer cells in TLS (Figure 6B), and this compartmental pattern was broadly reproduced across additional sections, although regional occupancy varied between sections (Supplementary Figure 6A–E). Both populations were detected inside and outside tumor regions (Figure 6C), establishing the shared anatomical context within which their interaction differences emerge.

**Figure 6.**
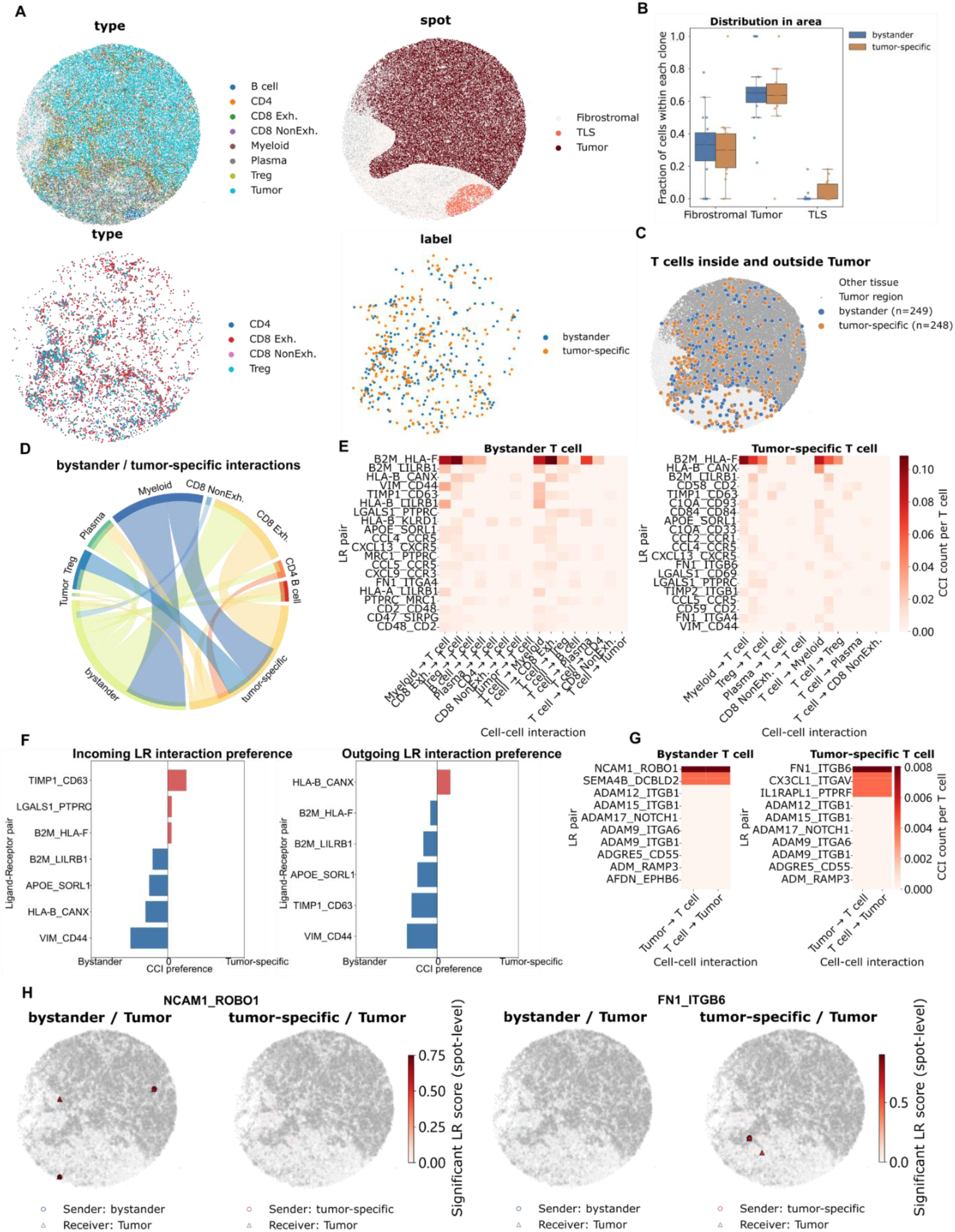
Spatial distribution and cell–cell interaction profiles of bystander and tumor-specific T cells in Slide-seq data. (A) Spatial maps colored by cell-type annotation, tissue region, T-cell subtype and BYSTER-predicted label. Tissue regions comprise fibrostromal areas, tumor areas and tertiary lymphoid structures (TLS). T-cell subtypes include CD4⁺ T cells, exhausted CD8⁺ T cells, non-exhausted CD8⁺ T cells and regulatory T cells. (B) Spatial distribution of bystander and tumor-specific clonotypes across the three tissue regions. For each clonotype, the fraction of its cells located within each region is shown. (C) Spatial distribution of bystander and tumor-specific T cells within and outside tumor regions. (D) Chord diagram summarizing inferred spatial cell–cell interactions (CCIs) between bystander or tumor-specific T cells and the annotated cell types. Ribbons represent interactions between the connected populations, with ribbon colors indicating the interacting cell populations. (E) Heatmaps of selected top LR pairs mediating inferred CCIs between the annotated cell types, including bystander and tumor-specific T cells. Color intensity represents the CCI count normalized per T cell. (F) Top significant LR pairs showing differential incoming and outgoing interaction preferences between the two predicted populations. Incoming interactions treat T cells as receivers, whereas outgoing interactions treat T cells as senders. Blue bars indicate preference for bystander cells, and red bars indicate preference for tumor-specific cells. (G) Heatmaps of selected top LR pairs mediating inferred interactions between tumor cells and each predicted T-cell population. Columns distinguish tumor-to-T-cell and T-cell-to-tumor interactions. Color intensity represents the CCI count normalized per T cell. (H) Spatial visualization of two representative LR pairs, NCAM1–ROBO1 and FN1–ITGB6, showing outgoing interactions from bystander or tumor-specific T cells to tumor cells. Circles indicate T-cell senders, colored blue for bystander and red for tumor-specific cells; triangles indicate tumor-cell receivers. Color intensity represents the significant LR score at the spot level.

**Supplementary Figure 6.**
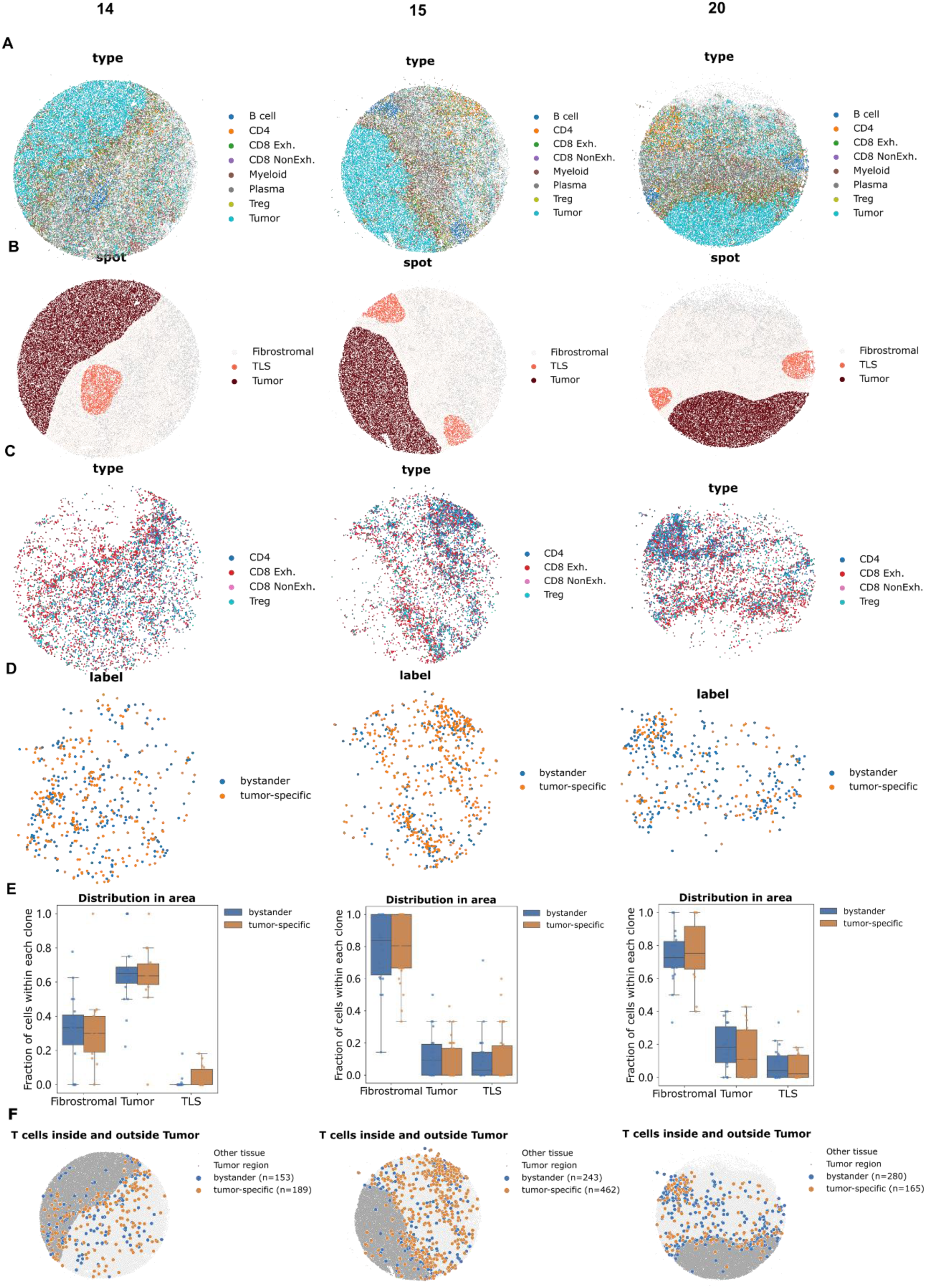
Spatial characterization of bystander and tumor-specific T cells in Slide-seq samples. (A) Spatial maps colored by cell-type annotation. (B) Spatial maps of fibrostromal regions, tumor regions and tertiary lymphoid structures (TLS). (C) Spatial distribution of CD4⁺, exhausted CD8⁺, non-exhausted CD8⁺ and regulatory T cells. (D) Spatial distribution of BYSTER-bystander and tumor-specific T cells. (E) Fractions of cells within each clonotype located in fibrostromal, tumor and TLS regions, grouped by bystander or tumor-specific identity. Each point represents a clonotype.

Spatial interaction analysis linked both predicted populations to myeloid cells, other lymphocytes and tumor cells (Figure 6D). Population-level ligand–receptor heatmaps revealed distinct communication programs in the two predicted clonal populations (Figure 6E). Although B2M/HLA-mediated axes dominated in both populations, bystander T cells additionally featured HLA-B–KLRD1 and CD47–SIRPG, whereas tumor-specific T cells preferentially engaged myeloid-derived signals such as C1QA–CD93 and CCL2–CCR1, with APOE–SORL1 represented in both. Differential preference analysis of shared pairs further uncovered direction-dependent roles. For both B2M–HLA-F and TIMP1–CD63, incoming interactions were preferentially assigned to tumor-specific T cells, whereas outgoing interactions were preferentially assigned to bystander T cells (Figure 6F), indicating that the two populations occupy distinct sender and receiver positions within shared communication axes.

Analysis restricted to communication with tumor cells further distinguished the predicted populations (Figure 6G). Among the displayed tumor-to-T-cell interactions, NCAM1–ROBO1 and SEMA4B–DCBLD2 were prominent for bystander cells, whereas tumor-specific cells showed prominent FN1–ITGB6, an extracellular matrix–integrin axis consistent with engagement with tumor stroma, and CX3CL1–ITGAV. Spatial mapping of outgoing T-cell-to-tumor interfaces for these two representative pairs localized the predicted T-cell senders relative to tumor-cell receivers and confirmed the compartment-specific detection of significant interfaces (Figure 6H), extending the transcriptional and clonal distinctions between bystander and tumor-specific T cells to the spatial dimension, where the two populations engaged in distinct, direction-dependent ligand–receptor communication within shared anatomical compartments.

## Discussion

Distinguishing bystander from tumor-specific T cells is a prerequisite for interpreting T-cell phenotypes in the tumor microenvironment. Bystander cells are abundant in human tumors and phenotypically distinct from tumor-reactive populations, as first demonstrated by Simoni et al. [10] in human colorectal cancer infiltrates and formalized in the framework of Meier et al.[2]. Yet neither transcriptional state nor TCR sequence alone provides a conclusive assignment: in the lung cancer cohort analyzed here, exhausted T-cell abundance, literature-derived specificity markers and clonal expansion each showed limited discrimination of clinical response. We therefore developed BYSTER, a heterogeneous graph neural network that combines scGPT-derived transcriptional representations with SCEPTR-derived paired TCR representations at metacell and metaTCR resolution, and propagates information across transcriptional, clonotypic and pathway-level relationships through cross-modal message passing.

BYSTER achieved consistent predictive performance across three datasets with distinct definitions of tumor specificity, with mean AUROCs of 0.856, 0.920 and 0.729 in the database-annotated corpus, experimentally labelled PDAC data and the lung cancer cohort, respectively, and outperformed unimodal and conventional multimodal baselines. Ablation analyses confirmed that this advantage derives from cross-modal message passing rather than simple feature concatenation, indicating that the relationship between T-cell state and antigen recognition is better represented through structured interactions between modalities than through their independent embeddings. The metacell and metaTCR framework further enables scalable analysis of large cohorts while retaining sample-specific relationships.

Beyond classification performance, the predicted populations revealed distinct biological programs associated with antigen-specific and antigen-independent T-cell activation. In NSCLC, bystander T cells were enriched for CD4⁺ naive and memory subsets, as well as MAIT cells, whose heterogeneity at single-cell resolution has been characterized by Garner et al. [30]. These cells also exhibited prominent inflammatory and stress-associated programs, including TNFα signaling via NF-κB and hypoxia. Increased JUN-family and NFKB1 regulon activity, accompanied by elevated expression of SLC4A10, LTK and CA2, further distinguished this population. In PDAC, bystander T cells exhibited relatively limited clonal expansion and enrichment of inflammatory and interferon-associated programs, whereas tumor-specific T cells showed greater clonal expansion under IP conditions and enrichment of proliferation-associated pathways. Notably, both cancer types showed concordant enrichment patterns, with cell cycle-associated programs, including E2F targets, G2M checkpoint and mitotic spindle, enriched in tumor-specific T cells, and inflammatory and stress-associated pathways, particularly TNFα signaling via NF-κB and hypoxia, enriched in bystander T cells. This cross-cancer concordance supports the reproducibility of these transcriptional differences across distinct tumor microenvironments and tumor-specificity annotation strategies. Collectively, these findings suggest that bystander T cells are not merely a passive residual compartment but a biologically distinct population potentially influenced by inflammatory cytokines and local microenvironmental cues. This interpretation is consistent with previous studies showing that inflammatory cytokines, including type I interferons and IL-15, can induce bystander activation of memory CD8⁺ T cells [11]. Furthermore, Leem et al. [31] demonstrated that IL-15-activated bystander CD8⁺ T cells contribute to tumor control in non-small cell lung cancer. Together, these observations suggest that cytokine-associated activation of bystander T cells may contribute to local immune regulation and influence the inflammatory context in which tumor-specific immune responses occur.

The tumor-specific T cells predicted by BYSTER were enriched for tissue-resident memory and exhausted states, consistent with previous work linking tumor antigen reactivity to these T-cell states, including the tissue-resident memory T-cell framework described by Amsen et al. [32] and the findings of Burn et al. [33]. However, our findings suggest that the relationship between T-cell identity and clinical response extends beyond the enrichment of tumor-specific populations. Responder samples contributed a larger proportion of the predicted bystander compartment, whose inflammatory and relatively non-proliferative phenotype may reflect cytokine-mediated activation within an immune-responsive tumor microenvironment[34], potentially involving IFNγ-associated signaling[35, 36]. This observation carries two implications for immunotherapy. First, the bystander compartment may serve as a cellular indicator of an inflamed, immune-responsive microenvironment, capturing aspects of immune activation that conventional measures of T-cell exhaustion or clonal expansion do not fully reflect. Second, our results highlight the limitations of interpreting clonal expansion or TCR-based tracking alone as evidence of tumor reactivity, as expanded intratumoral clones are not necessarily tumor-specific. Rather than reflecting competing immune populations, bystander and tumor-specific T cells may represent distinct but complementary components of the tumor immune microenvironment. These findings highlight the potential value of incorporating predicted bystander T-cell identity alongside conventional measures of tumor-reactive immunity when investigating immunotherapy responses.

Spatial analyses provided an additional perspective on how these populations interact with their local environment. In lung cancer Slide-seq sections Bystander and tumor-specific T cells inferred ligand– receptor interaction profiles differed. Bystander T cells showed interaction patterns associated with inflammatory communication and myeloid populations, whereas tumor-specific T cells displayed stronger associations with tumor-related extracellular matrix and integrin interactions. Spatial mapping of representative ligand–receptor pairs further illustrated the heterogeneous localization of these inferred interactions. These findings suggest that T cells occupying similar anatomical compartments can participate in distinct communication programs, linking predicted antigen specificity to the organization of local cellular interactions.

Several limitations should be noted. Our predictions are supervised by existing definitions of tumor specificity, and the lower performance in the lung cancer cohort indicates that database-derived annotations incompletely capture tumor reactivity in clinical samples. The association between the predicted bystander compartment and immunotherapy response is retrospective and requires validation in prospective cohorts, and the inferred ligand–receptor interactions, while spatially resolved, remain computational predictions rather than measured interactions.

Overall, BYSTER establishes a framework for integrating transcriptional and paired TCR information through biologically structured graph representations, with conserved bystander and tumor-specific programs across NSCLC and PDAC supporting its generalizability. By connecting predicted antigen specificity with cellular composition, clonal architecture, differentiation programs, clinical response and spatial communication, this approach provides a more comprehensive view of T-cell heterogeneity in cancer. The distinct yet overlapping features of bystander and tumor-specific T cells highlight antigen specificity as one component of a broader cellular identity shaped by receptor recognition, differentiation history and microenvironmental signaling. Incorporating these complementary dimensions can improve the interpretation of antitumor immunity and support more precise characterization of T-cell states in human tumors. From a translational perspective, BYSTER-derived bystander T-cell profiles may complement existing T-cell signatures for predicting immunotherapy response, warranting further validation in independent and prospective treatment cohorts.

## Declarations

### Ethics approval and consent to participate

#### Availability of data and material

All data and code needed to evaluate the conclusions in the paper are present in the paper and/or the Supplementary Materials. The code can be found at https://github.com/C8H8O-tenes/BTC

#### Funding

This work was partially supported by National Institutes of Health [R01LM014156, R01CA241930, R01GM153822, R01AA032723 to X.Z.]; National Science Foundation [2217515, 2326879 to X.Z.]; Cancer Prevention and Research Institute of Texas [RP250043 to X.Z.]; Center of Excellence — International Collaboration Initiative Grant (139170052).

#### Authors’ contributions

HX: conceptualization, data curation, formal analysis, investigation, methodology, project administration, writing–original draft.

JW: conceptualization, data curation, investigation, methodology, project administration.

KX: data curation, investigation, methodology.

XZ: conceptualization, funding acquisition, project administration

## Acknowledgements

We would like to thank all colleagues from our bioinformatics lab who provided their gracious support and advice

